# Evolution of a core ribosomal innovation in octopus

**DOI:** 10.64898/2026.06.25.734654

**Authors:** Rishav Mitra, Richard Han, Trey J Scott, Anik G Grearson, Jessica A Willi, Christina G Liu, Hyeongju Kim, Michael C. Jewett, Nicholas W Bellono, Amy SY Lee

## Abstract

Much of biology focuses on how genetic changes mediate new functions, but less attention is given to adaptations in other steps of the central dogma. Octopuses exhibit complex nervous systems and sophisticated behaviors that rival vertebrates, but via an entirely divergent evolutionary history. Here, we serendipitously discovered that octopus ribosomes contain a structural break in the core ribosomal RNA that is unique among all animals. This break site enhances translation fidelity to reduce miscoding and subsequent protein aggregation, even when engineered into evolutionarily distant bacterial ribosomes. Furthermore, high fidelity translation by octopus ribosomes supports proteomic stability during extensive RNA editing observed in cephalopods, suggesting synergy between distinct non-canonical modes of gene regulation. This adaptation emerged in recently derived octopuses with expanded nervous systems, thereby revealing a mechanism that could broadly support the evolution of novel organismal traits.

## Introduction

Octopuses carry out sophisticated behaviors that are comparable to those of complex vertebrates despite being evolutionarily distant relatives. In contrast to other invertebrates, octopuses exhibit a massively expanded nervous system, with neuron numbers similar to small primates ^1^. It remains unclear how such complex adaptations arose over the course of evolution, since octopuses do not exhibit vast genomic novelty when compared to other invertebrates ^2^. Post-transcriptional regulation of proteomic diversity has been proposed as a mechanism contributing to the elaboration of the cephalopod nervous system versus those of molluscan relatives ^3^; however, mechanisms of protein synthesis in octopus are virtually unexplored.

How do innovations in essential steps of the central dogma downstream of DNA sequence mediate novel functions? Ribosomes are the critical molecular machines that synthesize proteins to form the building blocks of all life ^4^. All ribosomes contain a common core of ribosomal RNAs (rRNAs) and proteins that direct protein synthesis. The rRNAs that make up the active sites for tRNA binding, mRNA decoding, and peptide bond catalysis are among the most evolutionarily conserved RNA sequences ^5^. Given this universal conservation, structural variation in the core architecture of the ribosome is rarely seen in nature ^6^. Therefore, how variations within the ribosome core facilitate changes in the fundamental process of protein synthesis remains poorly understood.

Here, we report the unexpected discovery that octopuses possess structurally unique ribosomes that enhance the accuracy of protein synthesis compared with ribosomes from all other tested animals and in heterologous chimeric systems. Enhanced translation fidelity synergizes with cephalopod-specific RNA hyper-editing to reduce protein aggregation. Furthermore, comparative analyses across octopus suborders and cephalopod relatives reveal that the ribosomal adaptation evolved in the clade of octopuses with expanded nervous systems. Thus, our findings suggest how adaptations within the core translation machinery could help drive the evolution of unique organismal traits and behaviors.

## Results

### The octopus has a unique structural adaptation in the conserved ribosomal core

During routine RNA extraction from octopus tissues (*Octopus bimaculoides*, **Fig. 1A**), we observed that the electrophoretic profile of 28S rRNA was split into distinct fragments (**Fig. 1B**). We further analyzed the rRNA using RT-PCR and RACE-PCR to reveal an octopus-specific sequence “break” localized between nucleotides G4322/C4323 in helix 88 (H88) of the E-site (**Fig. 1C**). This was particularly surprising because this is a highly conserved site in the ribosome that facilitates tRNA exit following peptide bond formation ^7^. Notably, this H88 rRNA cleavage is distinct from previously characterized breaks found in the highly variable D6 divergent regions located away from the ribosome catalytic core, such as those observed in protostomes ^8^, the naked mole rat ^9^, and tuco-tuco ^10^ (**Fig. 1C**). Sequencing confirmed a contiguous genomic rDNA locus, indicating that the H88 break site originates from a post-transcriptional cleavage event that gives rise to separate RNA fragments held together through non-covalent interactions (**Fig. 1B, C, Fig. S1A**). The break site was only found in octopus and not in any of the other diverse animals we analyzed with RACE-PCR (**Fig. 1D**). Furthermore, it was present across octopus tissues and developmental stages (**Fig. S1B**). Single-particle cryogenic electron microscopy (cryo-EM) structural analysis determined that the general architecture of the octopus ribosome was conserved compared with other organisms (overall RMSD=1.307Å; **Fig. 1E, Fig. S2, S3; Table S1**). Consistent with increased flexibility and the presence of an rRNA break, we could not resolve nucleotides 4321-4323 and observed a discontinuation in the rRNA density (**Fig. 1E**). Closer investigation of the rRNA revealed additional restructuring of the H88 loop compared to cryo-EM maps of mammalian ribosomes. In both octopus and human ribosomes, nucleotides in the loop of H88 base pair with six nucleotides in helix 22 of the 28S rRNA (octopus 28S rRNA nucleotides 324-329; human 28S rRNA nucleotides 317-322). However, while consecutive bases in the human H88 loop (nucleotides 4355-4360) fulfill these interactions, the equivalent base-pairing interactions in the octopus loop are distinct (**Fig. 1F**). On the 5′ side of the break, nucleotides 4314-4318 in the H88 loop base-pair with nucleotides 329-325 in helix 22, while on the opposite side, only U4324 base-pairs with A324. This structural alteration leads to an insertion of five nucleotides surrounding the break site (nucleotides 4318-4324), with U4320 stacking with A3537 in helix 68 of 28S rRNA, an interaction not observed in the human ribosome.

**Figure 1.**
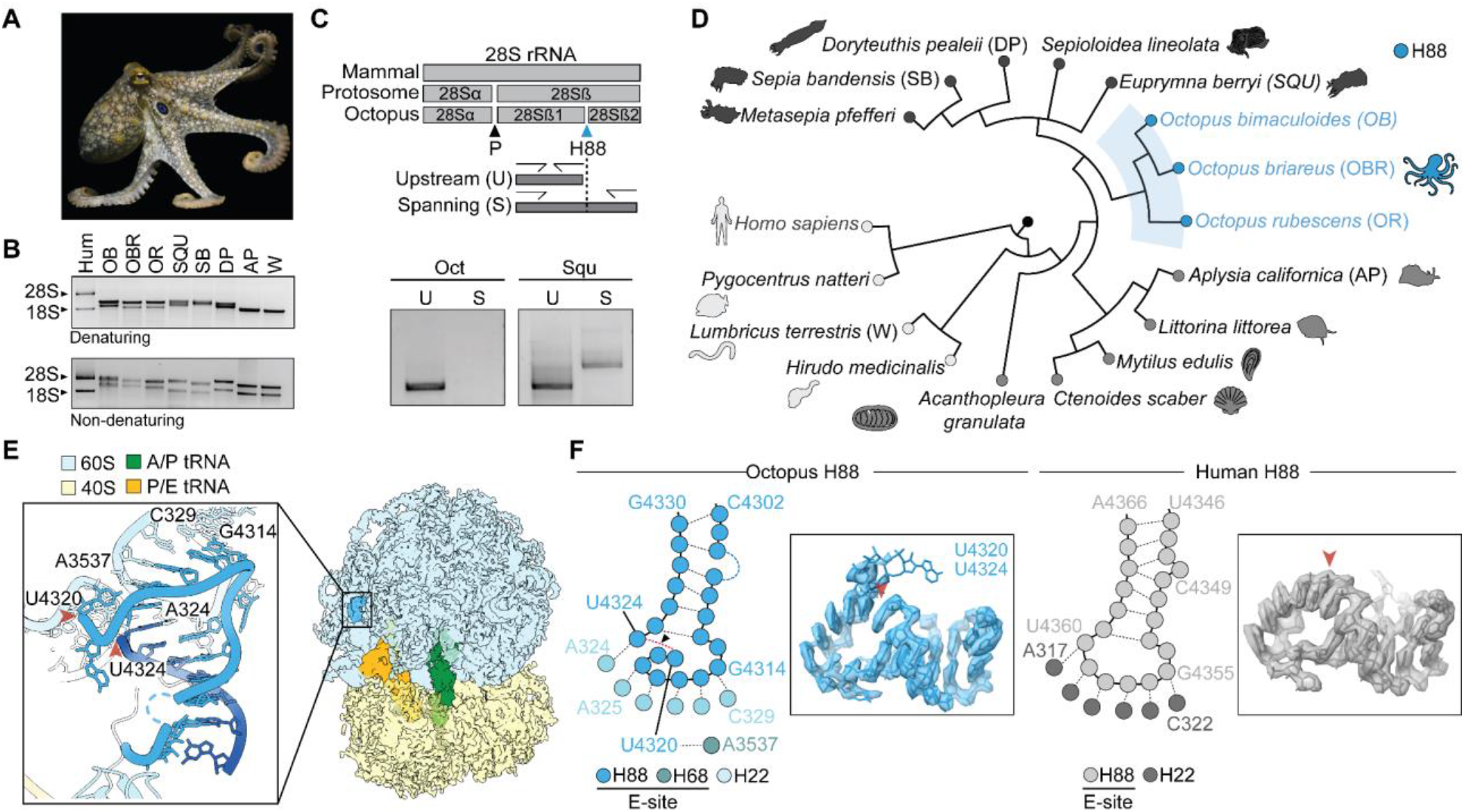
The octopus has a unique structural adaptation in the conserved ribosomal core. **(A)** California two-spot octopus (*Octopus bimaculoides*). **(B)** Denaturing and non-denaturing gel electrophoresis of total RNA revealed distinct sizes of rRNA fragments from human (HEK293T cells), *O. bimaculoides* (OB), *Octopus briareus* (OBR), *Octopus rubescens* (OR), *Euprymna berryi* (SQU), *Sepia bandensis* (SB), *Doryteuthis pealeii* (DP), *Aplysia californica* (AP), and *Lumbricus terrestris* (W). **(C)** Schematic of the unique octopus 28S rRNA break site in helix 88 (H88) compared to the conserved protostome rRNA gap (P). The octopus break was additionally validated by break-spanning RT-PCR. **(D)** Rapid Amplification of cDNA Ends (RACE)-PCR analysis of RNA isolated from phylogenetically diverse species including octopuses, cephalopods, mollusks, and others showed that the H88 break is unique to octopuses. **(E)** Cryo-EM structure of octopus ribosome in hybrid-PRE state resolved at 3.2 Å, with H88 highlighted (inset) **(F)** Predicted secondary RNA structure of H88, and the cryo-EM structures of octopus and human (PDB 6OLE ^37^) ribosomes highlighting a loss of electron density and altered base pairing interactions at the H88 break site. RNA numbering based off NCBI gene entry RNA28SN5.

### Octopus ribosomes have structural adaptations that facilitate accurate protein synthesis

The ribosomal E-site, including helix H88, contacts the 3′ end of deacylated tRNAs and is mechanically coupled to the L1 stalk network, subunit movements during translocation, and participates in allosteric communication with the decoding center ^11,12^. Notably, mutations in the A-site (H89-93) that decrease tRNA affinity by increasing local flexibility induce structural changes in helix 88, suggesting reciprocal communication between the A-site and E-site helices ^7,13^. In agreement, dimethyl sulfate (DMS) probing revealed heightened reactivity in the A-site of the octopus compared to the human ribosome, including in H89 and H91–93, which are part of the aminoacyl-tRNA accommodation corridor and important for selection of the correct tRNA during decoding ^14^ (**Fig. 2A–C**). These results indicate that octopus ribosomes exhibit increased A-site flexibility which could modulate key A-site functions such as tRNA binding.

**Figure 2.**
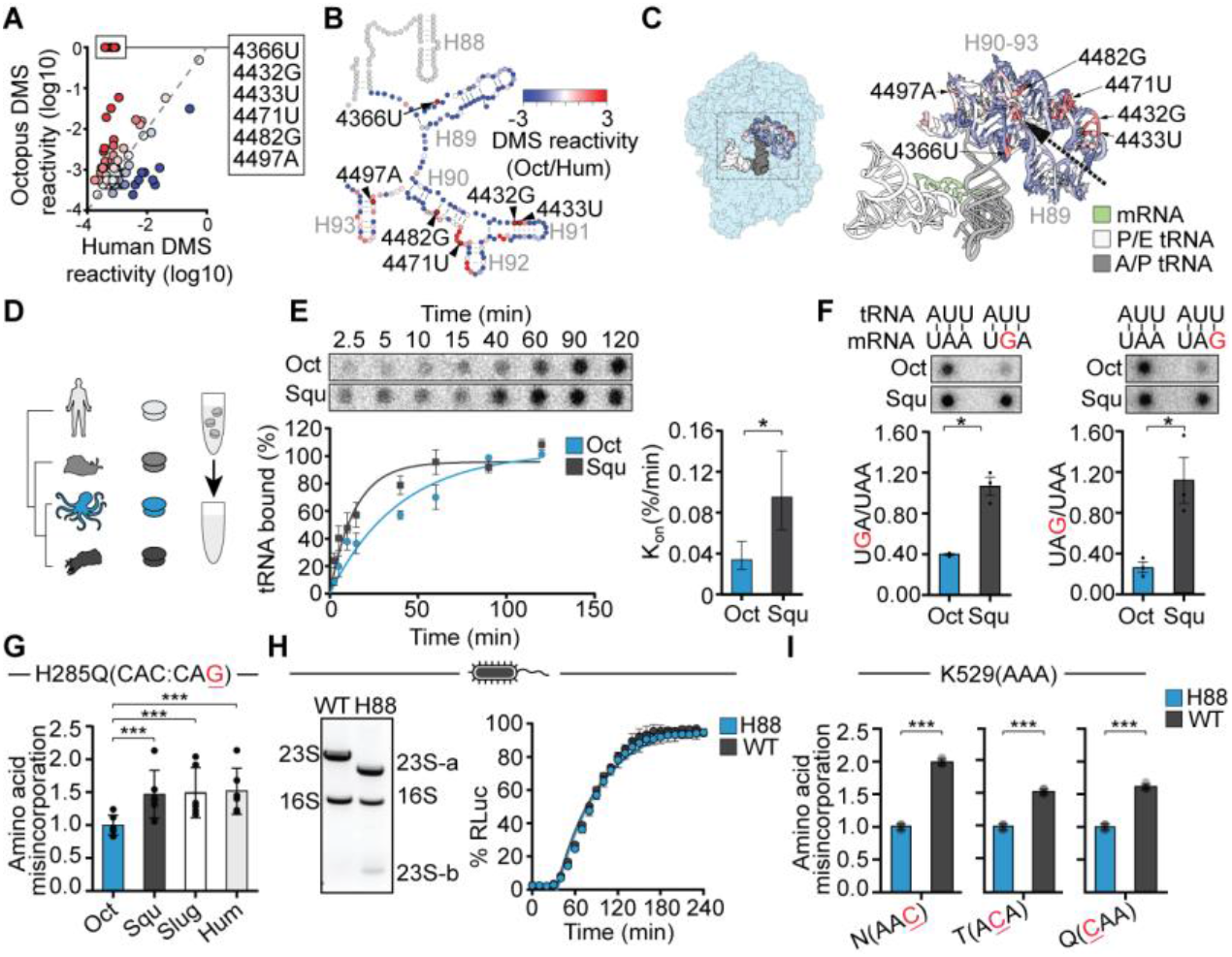
Octopus ribosomes mediate high-fidelity translation. **(A)** Comparative DMS probing of A-site nucleotides across human and octopus ribosomes revealed octopus-specific DMS-hypersensitive sites (4366U, 4432G, 4433U, 4471U, 4482G, 4497A). Residue numbers correspond to the human 60S structure PDB 6OLE^38^. **(B)** Nucleotides with increased flexibility in octopus ribosomes cluster in helices H89-93 within the A-site *aa-tRNA* accommodation corridor (black dashed arrow), as shown on the 2D rRNA and **(C)**, 3D ribosome structure (left) with the black dashed rectangle indicating the zoomed in region of interest (right). Residues have been color mapped as per log2 DMS reactivity (Oct/Hum) according to the scale in **(B). (D)** Ribosome-reconstituted *in vitro* reconstitution system to compare ribosomal function across species. **(E)** Kinetics of A-site tRNA binding in octopus (Oct) or squid (Squ) ribosomes using radiolabeled (^32^P) suppressor tRNA and ribosome-tRNA filter binding, with first-order rate constants (n = 5). **(F)** Octopus ribosomes exhibited more selective mRNA–tRNA pairing compared to squid ribosomes as measured by ribosome–tRNA filter binding using cognate and near-cognate codons (n = 3). **(G)** Octopus (Oct) ribosomes exhibited the highest translation fidelity compared with squid (Squ), *Aplysia* (*Slug*), and human (HEK293T; Hum) ribosomes as measured using a luciferase reporter for translation fidelity where amino acid misincorporation rescues luciferase activity (n=5). **(H)** *E. coli* ribosomes were reconstituted with the octopus H88 break, as seen by denaturing gel electrophoresis of total RNA (left) and *in vitro* translation kinetics (right). **I**, Chimeric ribosomes with the break decrease amino acid misincorporation rates across first, second, and third nucleotide codon positions (n = 6). Data represented as mean ± s.e.m. *p<0.05, **p<0.01, ***p<0.005 as determined by unpaired Welch’s t-tests.

To ask whether the structural features of the octopus ribosome correspond with functional differences, we developed an *in vitro* translation system that allows reductionist biochemical analyses of ribosomes isolated from non-model organisms. Briefly, we designed a ribosome-reconstitution system using ribosome-depleted rabbit reticulocyte lysates, ribosomes purified from wild caught animal tissues, and a luciferase mRNA reporter (**Fig. 2D, Fig. S4A, B)**. To perform comparative analysis of fundamental translation metrics, we chose octopus (*O. bimaculoides*) and diverse species without the H88 break, including the squid (*Euprymna berryi)* as another cephalopod, a marine slug (*Aplysia californica*) as a more distantly related mollusk, and human cells (HEK293T). The 28S rRNA across these species exhibits ~90% conservation in the 0.5 kb region flanking the octopus break site.

With this reductionist system, we first used radiolabeled (^32^P) suppressor tRNAs to quantify binding to the A-site (**Fig. S4C**). Octopus ribosomes exhibited overall lower tRNA binding affinity and, importantly, a 4-fold reduction in affinity for near-cognate tRNA–mRNA pairing compared to that of squid (**Fig. 2E, F**). Thus, octopus ribosomes exhibit increased decoding stringency and favor highly selective mRNA–tRNA pairing. In agreement with the enhanced decoding stringency, we next assessed translation accuracy using a luciferase-based reporter for translation error ^15^. Protein synthesis mediated by octopus ribosomes exhibited significantly increased accuracy compared with other species, accompanied by reduced translation efficiency (**Fig. 2G, Fig. S4B**).

To test if the octopus H88 break is sufficient to confer enhanced translation fidelity, we used a heterologous *in vitro* reconstituted ribosome engineering system from *E. coli* ^16^. Chimeric *E. coli* ribosomes engineered to contain the octopus H88 rRNA break showed no change in translation efficiency, but exhibited a 2-fold increase in fidelity (**Fig. 2H, I**).

Thus, the octopus-specific H88 ribosome adaptation is sufficient to confer enhanced accuracy of protein synthesis, even when transplanted into evolutionarily distant prokaryotic ribosomes. Notably, engineering similar increases in translation fidelity has broad effects on evolutionary fitness, indicating that the H88 break may represent a beneficial adaptation in octopus ^17^. Indeed, the 2-fold change in translation fidelity conferred by the H88 break is comparable to that achieved by ribosomal protein mutations that extend lifespan and stress resistance across diverse species^18^.

### Octopus ribosomes support reduced protein aggregation

What is the cellular impact of enhanced translation fidelity in octopus? Since mistranslation increases protein misfolding and aggregation ^19^, we first measured protein aggregation in native octopus tissues. We focused on the nervous system, which is elaborated in octopus and generally enriched in long-lived proteins highly sensitive to translation errors. Under homeostatic conditions, octopus axial nerve cords contained fewer and smaller aggregates compared to squid (**Fig. 3A, Fig. S5B**). To examine if this phenotype is sensitive to ribosomal decoding fidelity, we treated *ex vivo* tissue explants with G418, an aminoglycoside antibiotic that induces miscoding by binding to the ribosomal decoding center ^20^ (**Fig. 3B, Fig. S5A**). G418 treatment increased aggregate size in octopus nerve cords and partially reduced the difference in aggregation between species, shifting octopus tissues towards a more squid-like aggregation state (**Fig. 3B**). Thus, the low aggregation state of octopus tissue is sensitive to decoding perturbations, suggesting that ribosomal decoding stringency could contribute to reduced proteotoxic burden. Indeed, insoluble protein aggregates isolated from native tissues across cephalopod relatives and other mollusks demonstrated that octopuses had fewer aggregated proteins across multiple gene families (**Fig. 3C, D**).

**Figure 3.**
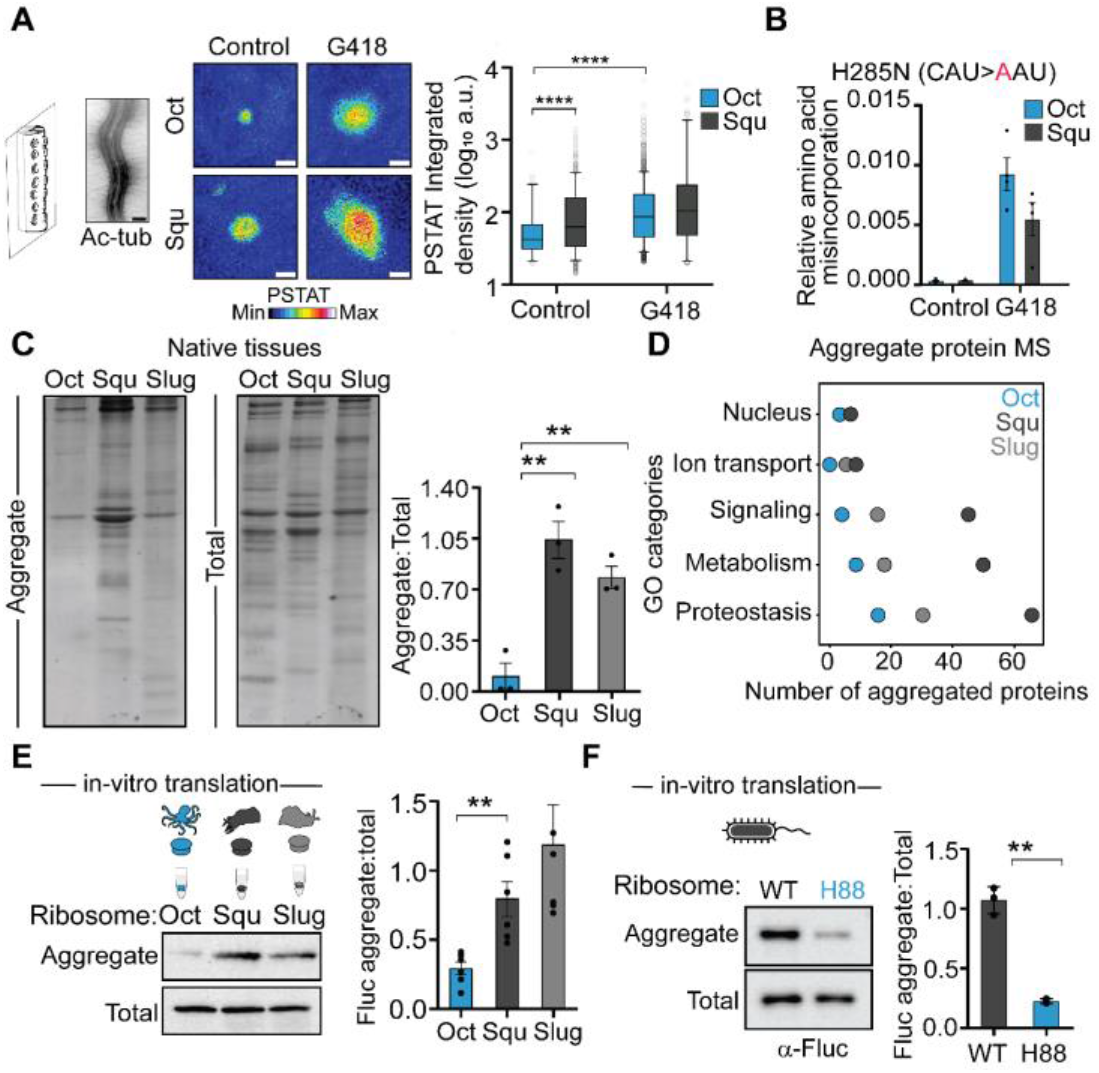
The H88 ribosomal break facilitates protein synthesis with reduced aggregation. **(A)** Octopus axial nerve cords (marked by acetylated α-tubulin; scale bar=500 µm) had smaller aggresomes compared with squid as visualized by comparative imaging of proteostat (PSTAT)-labeled protein aggresomes, inclusion bodies composed of aggregation-prone proteins. Pharmacological reduction of translation fidelity with G418 increased octopus neuronal aggregation. Scale bar = 1 µm. Data represented as mean ± s.e.m.; n = 4 animals across conditions. **B**, Pharmacological attenuation of translation fidelity with G418 increased misincorporation by octopus to levels comparable to untreated squid as measured using mRNA reporters for translation error (n = 4). **C**, Native tissue insoluble aggregates were lowest in octopus compared with squid and slug, as measured by SDS-PAGE and Coomassie staining of total versus insoluble proteins (n = 3). **D**, Gene ontology (GO) comparison of aggregated proteins derived from native tissues of octopus (Oct), squid (Squ) or *Aplysia* (Slug) using mass spectrometry showed no preferential enrichment in any particular category. **E**, Octopus ribosomes produced fewer aggregated proteins compared with squid and slug ribosomes as measured by *in vitro* translation of Fluc, aggregated protein separation by centrifugation, and immunoblotting (n = 6). **F**, Chimeric *E. coli* ribosomes with the octopus H88 break produced fewer aggregates when compared to WT ribosomes (n=3). Data represented as mean ± s.e.m. *p<0.05, **p<0.01, ***p<0.005 as determined by unpaired Welch’s t-tests.

To directly assess the role of the octopus ribosome in producing fewer protein aggregates, we first measured *in vitro* aggregation of Fluc translated by ribosomes from different species. Translation by octopus ribosomes produced Fluc protein with reduced aggregation propensity compared to squid or slug ribosomes (**Fig. 3E**). Importantly, chimeric *E. coli* ribosomes engineered to contain the octopus H88 rRNA break also generated Fluc protein with lower aggregation compared to protein synthesized by wild type ribosomes (**Fig. 3F**). Thus, the octopus H88 break directly enhances proteostasis. Consistent with this functional outcome, accurate protein synthesis in octopus is paired with reduced proteasome activity, lower expression of proteostatic machinery, associated signaling pathways, and relaxed evolutionary selection of proteasomal genes (**Fig. S5C–H**). Collectively, these findings indicate that octopus ribosomal innovation supports global proteostasis.

### Octopus ribosomes promote accurate decoding of edited RNA

While the central dogma describes the flow of genetic information from DNA to RNA to protein, organisms can bypass this strict conservation of genomic sequence through post-transcriptional modifications that alter mRNAs. In coleoid cephalopods, including octopus and squid, this occurs through a form of post-transcriptional modification called hyperactive adenosine-to-inosine (A-toI) mRNA editing ^3^. RNA editing is particularly prevalent in the expanded cephalopod nervous system, with octopuses containing between 80-130,000 sites in protein coding regions that are subject to adaptive RNA editing and result in recoded proteins ^3,21,22^ (**Fig. 4A**). However, in other organisms, mRNA editing can lead to nonsynonymous substitutions and misfolding, ribosome stalling, or truncated proteins ^23,24^. Considering the octopus ribosome exhibits more stringent mRNA–tRNA pairing (**Fig. 2E**), we hypothesized that this function could facilitate selective decoding of inosine-containing mRNAs. To characterize how octopus ribosomes interpret edited mRNAs during the decoding process, we examined base pairing by measuring tRNA binding to mRNAs modified to contain inosine at a single codon position. Consistent with highly selective tRNA pairing, octopus ribosomes showed the highest binding affinity for the most thermodynamically favored inosine-cytosine pair and lower affinity for other promiscuous pairs (**Fig. 4B**). This was in contrast with ribosomes from squid, which exhibited more promiscuous tRNA binding, or human ribosomes which stall at inosines and cannot translate highly edited mRNA ^25^ (**Fig. 4B**). Notably, both cephalopod ribosomes were able to translate an inosine-containing FLuc reporter, in contrast to human ribosomes which likely stall at inosines ^26^. Consistent with promiscuous tRNA-mRNA pairing, translation of edited mRNAs by squid ribosomes resulted in a substantial increase in aggregation compared with translation of unedited mRNAs (**Fig. 3E and Fig. 4C, D**). However, octopus ribosome-mediated translation resulted in 10-fold lower protein aggregation compared to squid (**Fig. 4C, D**). While reduced aggregation of inosine-containing mRNA by octopus ribosomes recapitulates the stringent decoding conferred by the H88 break in heterologous systems (**Fig. 3F**), other native ribosomal properties could contribute. Together, these findings suggest that the ability of octopus ribosomes to selectively decode inosine-containing mRNAs could synergize with RNA editing while limiting misfolding and aggregation.

**Figure 4.**
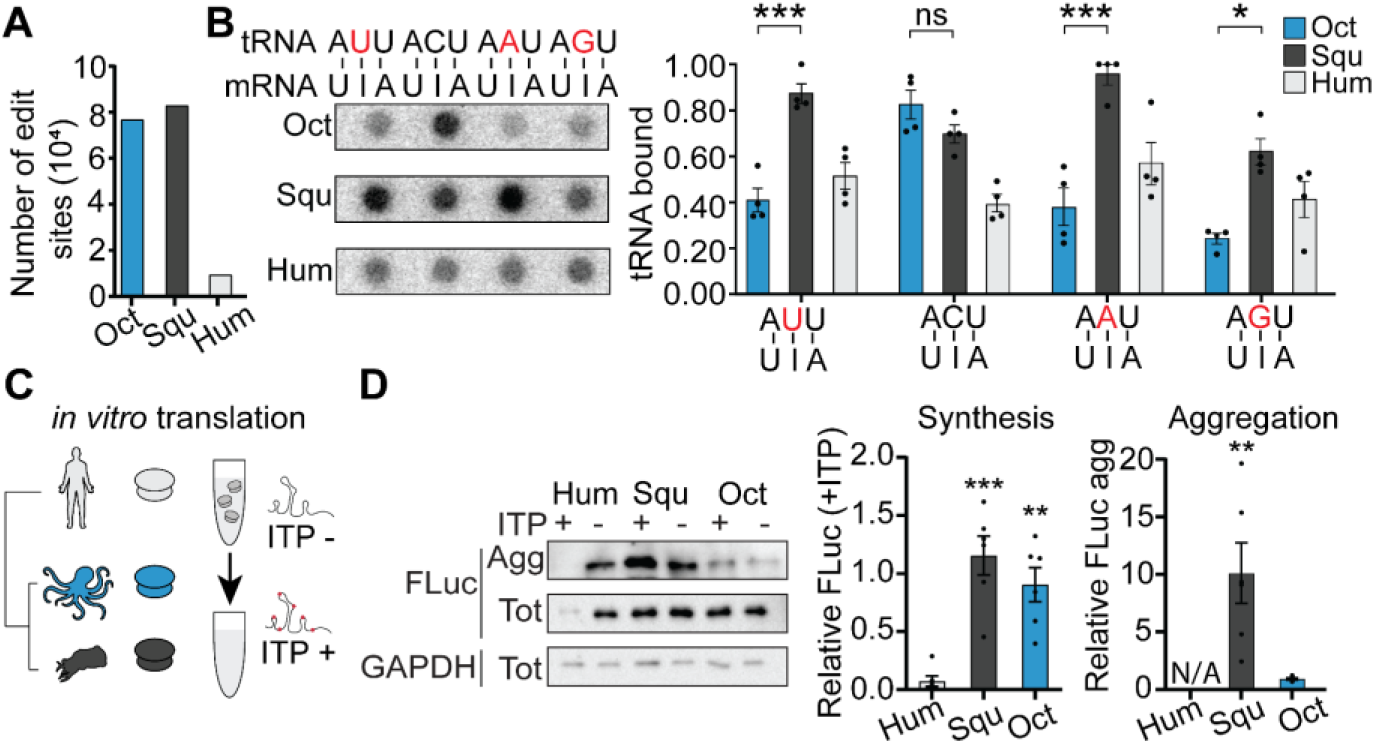
Octopus ribosomes translate hyper-edited transcripts with reduced aggregation. **(A)** Octopus (Oct) and squid (Squ) perform extensive adenosine-to-inosine (A-to-I) RNA editing compared with humans (Hum), at a greater number of editing sites. **(B)** Octopus ribosomes selectively decode inosine-containing mRNA codons (strongest I:C pairing) whereas squid ribosomes show promiscuous tRNA binding, as measured by ribosome-tRNA filter binding using an mRNA containing inosine (n = 4). Inosine has the highest binding affinity for cytosine (ΔG= −8.8 Kcal/mol), but can also weakly bind adenosine (ΔG= −7.5 Kcal/mol), uracil (ΔG= −5.9 Kcal/mol), or guanosine (ΔG= −6.3 Kcal/mol)^39^. **(C-D)** Octopus and squid, but not human, ribosomes can translate inosine-containing firefly luciferase (*Fluc*) reporter mRNA. Octopus ribosomes produced less aggregated protein than squid ribosomes when translating edited mRNAs. Proteins produced by squid ribosome-mediated translation of edited mRNAs showed 10-fold higher aggregation versus unedited mRNAs (2.6-fold, see Figure 3E), while octopus ribosomes showed no effect of RNA editing on aggregation. ITP = *in vitro* transcribed mRNAs containing inosine, n = 6. Data represented as mean ± s.e.m.; *p<0.05, **p<0.01, ***p<0.005, ****p<0.0001 as determined by unpaired Welch’s t-tests.

### Evolution of a lineage-specific ribosome innovation

The functional sites (A, P, E) of the ribosome are highly conserved across all domains of life, and rRNA breaks outside of the octopus are restricted to highly divergent surface-exposed regions away from the core center ^8-10^. How, then, did the H88 break originate? Octopus and squid are separated by almost 300 million years of evolution^27^. Thus, we sought to analyze the H88 rRNA break across more closely related lineages to understand the evolutionary origins of the break. Octopuses comprise two major suborders: 1) Incirrates, the most prevalent octopuses that live within shallow depths and utilize a highly expanded nervous system to thrive within variable, predator-rich environments using their autonomous sensory arms for hunting, complex camouflage behaviors, and the capacity for learning and rapid decision-making ^28,29^; and 2) the understudied and often inaccessible Cirrates, which diverged over 100 million years ago in the Late Cretaceous period ^30^ and inhabit deep-sea environments with minimal sensory input by using simpler nervous systems to facilitate slow-swimming and passive-feeding lifestyles ^31,32^ (**Fig. 5A, B; Fig. S6A**). To test whether the H88 rRNA break evolved in concert with nervous system expansion in Incirrates, we analyzed a deep sea Cirrate (*Grimpoteuthis*) collected at a depth of 406m via slurp-sampling by a remotely operated vehicle (**Fig. 5C**). Strikingly, comparative RT-PCR of RNA extracted from *Grimpoteuthis* tissue demonstrated complete absence of the H88 rRNA break that we observed in every other Incirrate octopus examined (**Fig. 5D, Fig. 1D**). To more comprehensively characterize the extent of the H88 rRNA break in cephalopods, we searched for RNA sequence determinants across cephalopods. The rRNA break occurs at a highly conserved region of the large ribosomal subunit that emerged early in ribosomal evolution ^33^. Remarkably, the break site corresponds to a specific DNA sequence (TATG/CGTC) that has been perfectly conserved across almost 100 million years of diversification in the Incirrata lineage of cephalopods (**Fig. 5E**). This conserved break site sequence represents a lineage-specific innovation, with Incirrates showing numerous substitutions around the break, despite all other cephalopods and other animals exhibiting high conservation in this region (**Fig. S6C, D**). Intriguingly, this divergent evolutionary path is mirrored in the ribosomal break site and in the distinct differences in nervous system expansion across these two clades of octopuses. (**Fig. 5A, E, S6A–D**).

**Figure 5.**
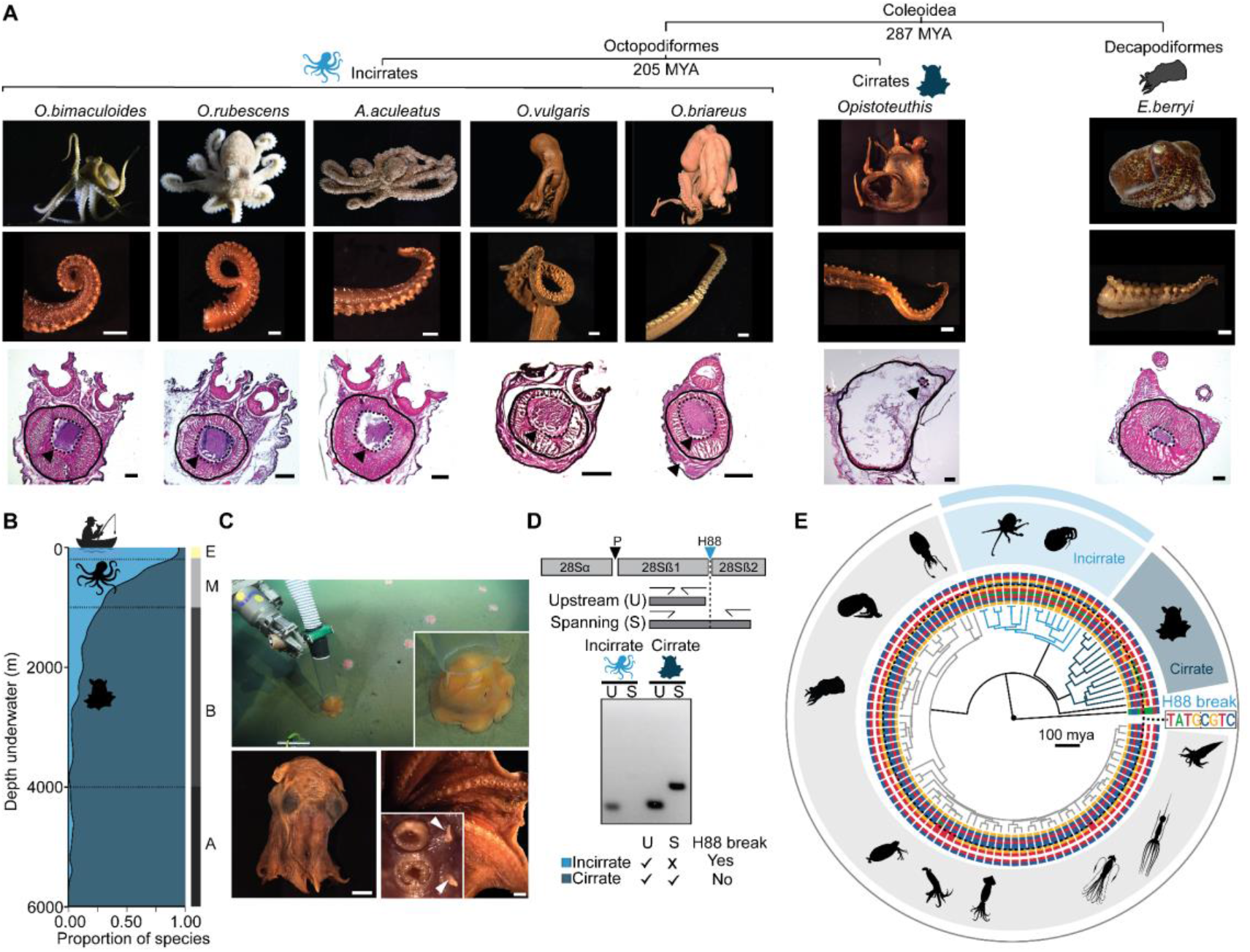
The octopus H88 break is a lineage specific innovation that evolved in Incirrate octopuses. **(A)** Phylogenetic tree for coleoid cephalopods with comparative arm histology of Incirrate (*Octopus bimaculoides, Octopus rubescens, Abdopus aculeatus, Octopus vulgaris, Octopus briareus*), Cirrate (*Opistoteuthis*) and Decapodiformes (*Euprymna berryi*). Arm cross sections reveal an expanded axial nerve cord in the Incirrates. Scale bar=1mm for arm images and 500 µm for arm cross sections. **(B)** Depth distributions indicate that Incirrates inhabit shallow waters while their closest relatives, the Cirrates are predominantly found in the deep sea. The right bar indicates zone layers of the ocean: E, Epipelagic (yellow); M, Mesopelagic (light grey); B, Bathypelagic (dark grey); A, Abyssopelagic (black). **(C)** Collection of a deep sea Cirrate octopus (*Grimpoteuthis*) using a remotely operated vehicle (ROV), top. Bottom, museum specimens of Cirrate octopuses (*Opistoteuthis*). Inset: magnified view of arms and sucker cups. Scale bars (left to right) = 1 cm, 500 µm, 2 mm. White arrows point to the cirri. **(D)** rRNA RT-PCR spanning the H88 region reveals that the H88 break site is unique to Incirrates. **(E)** The H88 break site is highly conserved in only the Incirrate lineage of octopuses (light blue) but absent in the Cirrates (dark blue) other cephalopods (grey). Comparative phylogenetic analysis across cephalopods shows that the break sequence is an Incirrate-specific innovation in a region of high genomic conservation.

## Discussion

The ribosome contains highly conserved foundational structures that trace back billions of years, especially within the core rRNA regions that form the A-, P- and E-sites involved in mRNA decoding and peptide bond synthesis. Here, we show that a naturally occurring rRNA break in E-site of the octopus ribosome modulates central ribosome function to enhance translation fidelity and reduce protein aggregation. Importantly, engineering the octopus rRNA break into a highly divergent prokaryotic ribosome confers higher translation fidelity, highlighting the evolutionary modularity of the ribosome to accommodate adaptations while preserving essential catalytic function. Indeed, single amino acid substitutions in ribosomal proteins distant from the core of the ribosome have been shown to enhance translational fidelity and extend lifespan and stress resistance in yeast, worms, and flies ^18^. While octopus ribosomes translate ~30% slower than those from squid or slug, this property is independent of the H88 break, which is sufficient to enhance fidelity without slowing translation rates in *E. coli*. Thus, the H88 break decouples accuracy gains from canonical speed-fidelity trade-offs. Altogether, these findings suggest that ribosome innovations could support specific organismal demands in a manner that is more widespread than currently appreciated, and therefore represents a key aspect of the central dogma to be further explored.

Unlike vertebrates, nervous system expansion in octopuses appears to be driven by ecological rather than social factors ^34^, as their sophisticated behaviors evolved in largely solitary and short-lived lifestyles ^35,36^. Because Incirrate octopuses require rapid gene expression regulation to survive in a dynamic shallow marine environment, we speculate that high-fidelity translation may have been favored as it supports proteomic flexibility through RNA editing. Squid, lacking this ribosomal innovation, instead evolved hyperactive proteasomes (**Fig. S5C**–**H**), which could mitigate the proteotoxic burden of RNA editing. Altogether, octopuses exhibit innovations across levels of the central dogma. These synergistic adaptations point to a distinct evolutionary strategy that allows for enhanced phenotypic flexibility.

### Limitations of the study

Here, we use *in vitro* biochemistry, reductionist systems, and phylogenetics to reveal how an rRNA break in the octopus ribosome alters ribosome function and proteostasis. However, a limitation of our study is the lack of experimental tools to study octopus protein synthesis *in vivo*, with nearly no protocols established in wild-caught animals outside of this work. Future development of challenging genetics and primary cell culture approaches would allow us to further probe the synergy between ribosome adaptations and gene regulation programs, and ultimately their relationships to complex animal physiology. Furthermore, direct comparison of ribosomes from Cirrate and Incirrate octopus relatives that differ in natural habitat and presence of the H88 rRNA break would be of great interest; however, obtaining deep-sea Cirrates is extraordinarily difficult and rare. While numerous technical challenges remain, our initial discovery of an innovation within octopus ribosomes establishes powerful conceptual approaches in comparative biochemistry and heterologous systems. Furthermore, the relationship between species-specific ribosomes and organismal behavior provides a foundational blueprint for studying ribosome evolution and function across the diversity of life.

## Supplemental Information

Document S1. Figures S1–S6 and Table S1.

## Acknowledgements

We thank C. Winkler for providing octopuses; B. Grasse of the Cephalopod Program and the Marine Resource Center at the MBL for squid and other invertebrates; B. Walsh and P. Kilian for assistance with animal care and experiments; the Harvard Center for Biological Imaging for imaging and consultation services; and the Taplin Mass Spectrometry Facility for proteomics services; J. Gao, M. Yip, S. Shao, the Harvard Center for Cryo-Electron Microscopy (HC2EM), and SBGrid for help with structure determination and analysis; P. Girguis and the Scripps Institution of Oceanography Benthic Invertebrate Collection (SIO-BIC M17639, W7150, M11294, M11232, M11058); the captain and crew of R/V *Falkor* and the pilots and technicians of ROV *SuBastian* (cruise number FK181005), members of the Bellono lab, Lee lab, and K. Chat for discussions; L. Soucy for the graphical abstract. This research was supported by grants to N.W.B. from the New York Stem Cell Foundation, the Harvard NeuroDiscovery Center, and the National Institutes of Health (R35GM142697); grants to A.S.Y.L from the Pew Biomedical Scholars Program, the Mathers Foundation, and the National Institutes of Health (R35GM142527); a Harvard Brain Science Initiative (HBI) Postdoctoral Pioneers grant to R.M.; an NRSA fellowship to R.H. (F31NS132412); T.J.S is a fellow of the Jane Coffin Childs Fund for Medical Research.

## Author contributions

Conceptualization, A.S.Y.L. and N.W.B.; Methodology, R.H., R.M., A.G.G., J.A.W., C.G.L.; Investigation, R.H., R.M., A.G.G., J.A.W., C.G.L., H.K.; Formal analysis, R.M., T.J.S.; Writing – original draft, R.M., A.S.Y.L., N.W.B.; Writing – review & editing, all authors; Supervision, A.S.Y.L. and N.W.B.; Funding acquisition, A.S.Y.L. and N.W.B.

## Data and Materials availability

Sequencing data are deposited at NCBI under BioProject PRJNA1199547 and GEO accession GSE284564. Atomic model coordinates and cryo-EM density maps are deposited at PDB under accession code 9EHO and EMDB under accession codes EMD-48049, EMD-48050, EMD-48051, and EMD-48061. *Lead contact:* Further information and requests for resources and reagents should be directed to and will be fulfilled by the lead contacts-Nicholas W. Bellono and Amy SY Lee.

