## Supplementary Materials for "Evolution of a core ribosomal innovation in octopus"

This PDF file includes

#### Supplementary Figures 1-6

#### Supplementary Table 1

#### Materials and Methods

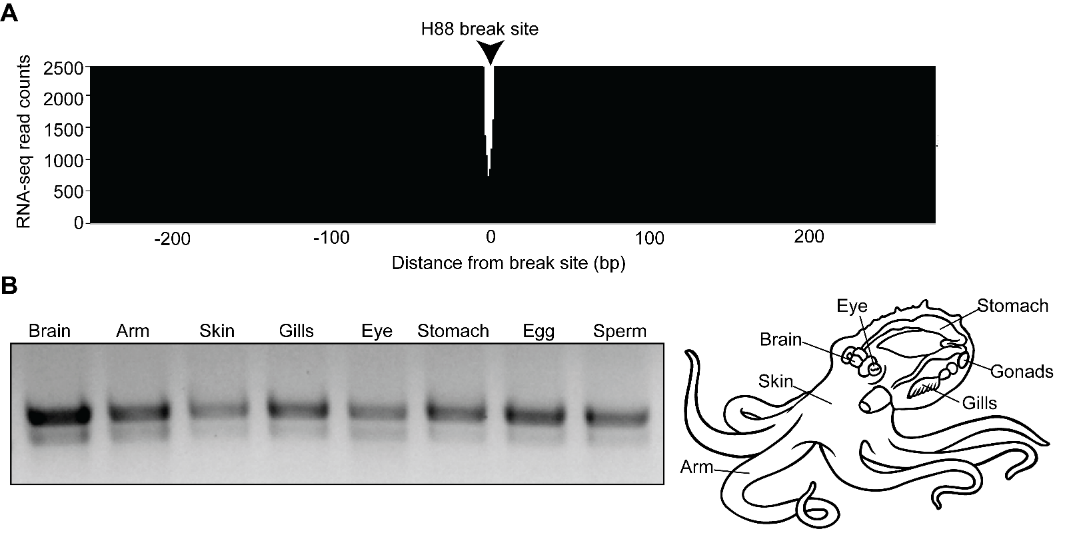

Figure S1: **Octopus ribosome adaptation across tissues**. (A) Amplicon sequencing from cDNA isolated from octopus ribosomes confirms a gap in reads spanning the H88 break site. (B) Denaturing gel electrophoresis of RNA isolated from *Octopus bimaculoides* tissues shows that the unique octopus rRNA break exists in all examined tissue types including brain, arm, skin, gills, eye, stomach, egg, and sperm.

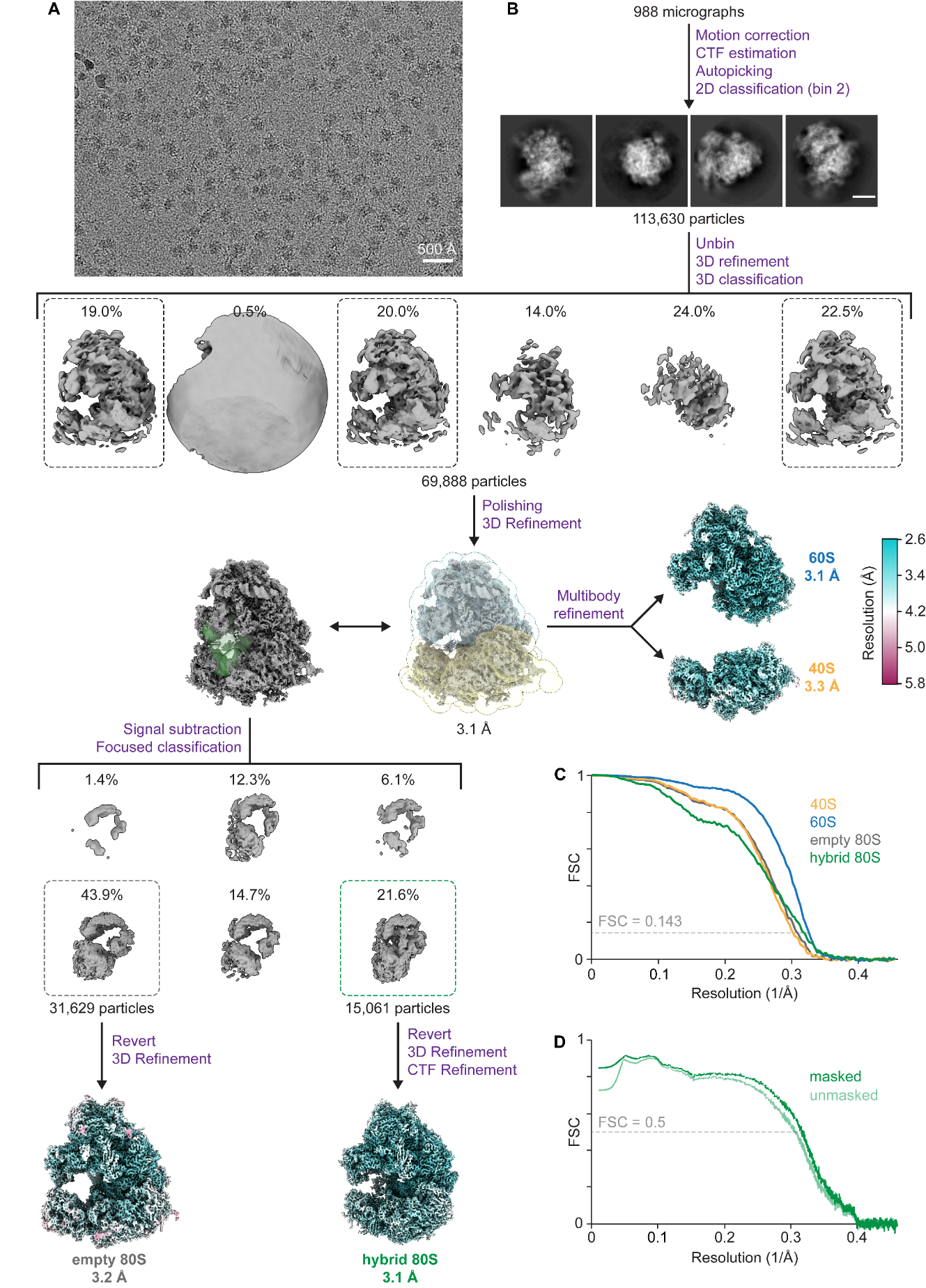

Figure S2: **Cryo-EM data processing.** (A) Representative cryo-EM micrograph of purified *Octopus bimaculoides* monosomes. (B) Cryo-EM data processing and classification strategy. Maps used for modelling are colored by local resolution. Masks used for multibody refinement (blue – 60S; yellow – 40S) and focused classification for tRNA occupancy (green) are indicated. Scale bar, 100 Å. (C-D) Fourier shell correlation (FSC) vs. resolution (1/Å) curves for the indicated cryo-EM maps.

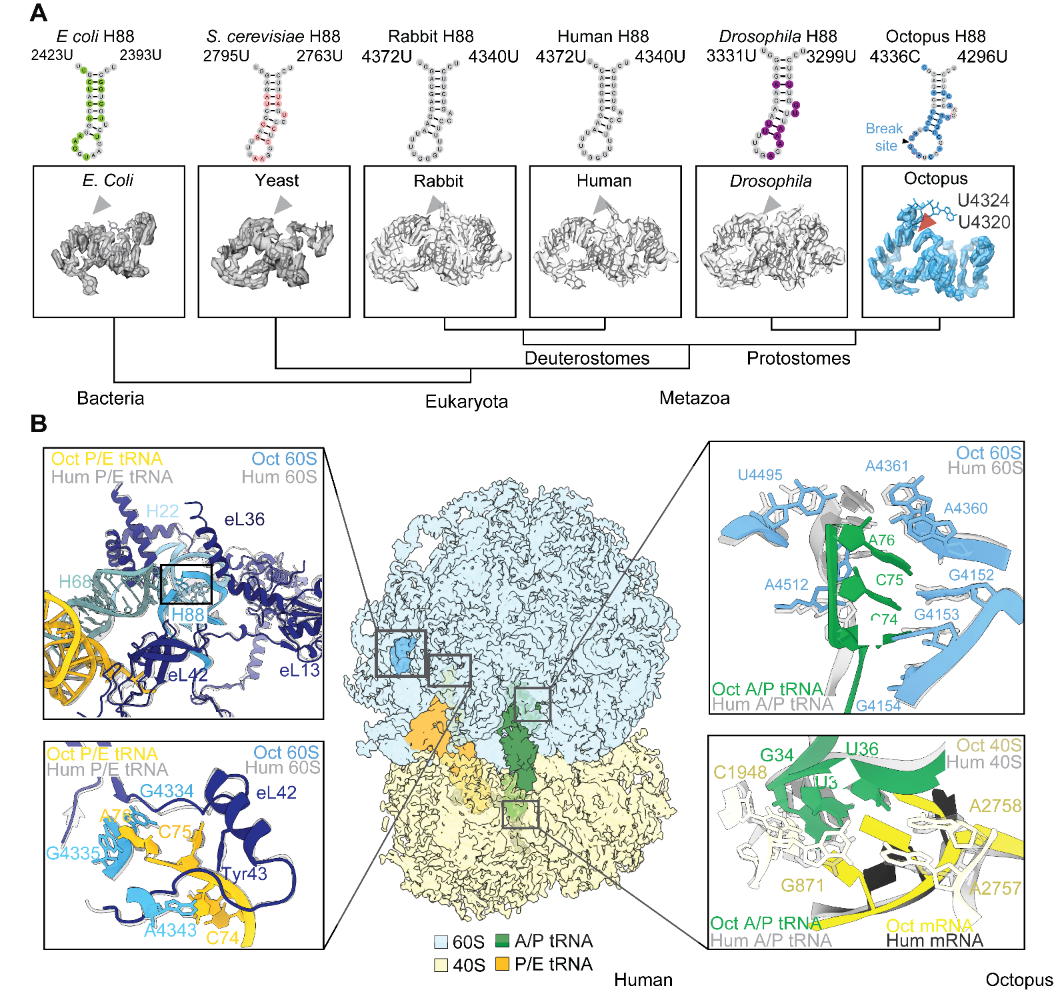

Figure S3: **Octopus ribosome adaptation across tissues, and structural comparisons.** (A) H88 structural conservation across species showing cryo-EM density, nucleotide residues, and coordinates overlaid with octopus H88 (Human: 6Y57^75^, Yeast: 8CMJ^77^, Rabbit: 6HCJ^76^, *Drosophila*: 4V6W^76^, *E. coli*: 7N2U^78^). (B) Octopus ribosome overlaid with the human ribosome (PDB 6Y57^5^) showing the location of the H88 break and shift in nearby protein eL13 (top). CCA ends and interacting region of P/E tRNA in octopus and human ribosomes (bottom). CCA ends and surrounding region of A/P tRNA (top). A/P tRNA anticodon and mRNA surrounded by proofreading bases in octopus and human ribosomes (bottom).

Figure S4: **Comparison of ribosome translation efficiency and tRNA binding affinity across species.** (A) Renilla Luciferase (RLuc) activity after 3 h in rabbit reticulocyte lysate (RRL) compared to ribosome-depleted RRL supplemented with no ribosomes (-) or ribosomes purified from octopus (Oct), squid (Squ) or *Aplysia* (Slug) tissue. (n=3) (B) Time course of luciferase activity during ribosome-reconstituted *in vitro* translation of *RLuc* mRNA by octopus, squid, or slug ribosomes. Protein synthesis is normalized to the relative maximum for calculation of first-order rate constants. (C) Schematic representation of ribosome–A-site tRNA binding assay using a radiolabeled (^32^P) suppressor tRNA, a start-stop codon mRNA (AUG–UAG), octopus or squid ribosomes, and measured by filter binding. Diphtheria toxin (dtA) is added to block eEF2 activity, ribosome translocation, and thereby examine A-site tRNA binding. Data represented as mean ± s.e.m.; *p<0.05, **p<0.01, ***p<0.005 as determined by unpaired Welch’s t-tests.

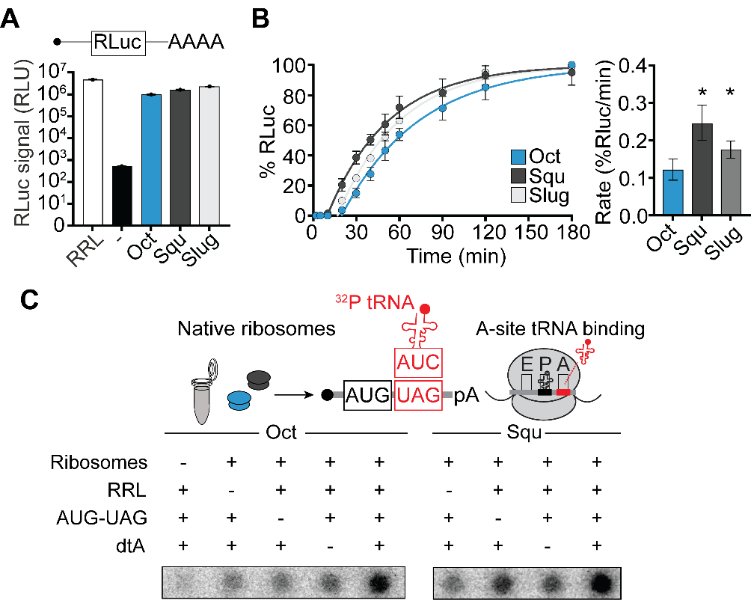

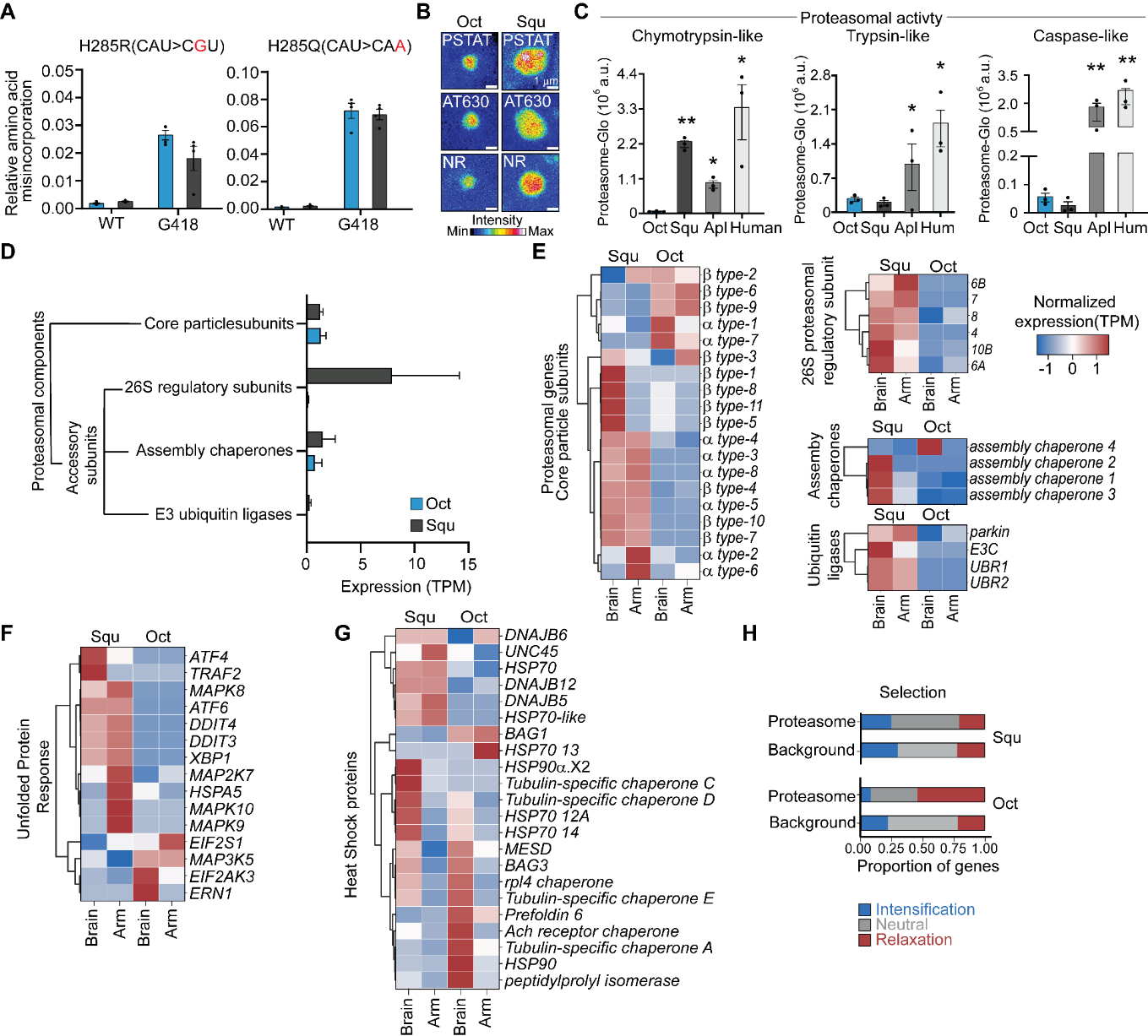
Figure S5: **Proteostasis in cephalopod tissues.** (A) Pharmacological reduction of translation fidelity with G418 increased misincorporation by octopus to match squid, slug, and human as measured using mRNA reporters for translation error (n = 4). (B) Comparative imaging of proteostat (PSTAT)-labeled protein aggresomes in octopus and squid central brains indicated that octopuses had smaller aggresomes compared to squid. Detection of aggresomes in octopus or squid axial nerve cords was consistent across Proteostat (PSTAT), Amytracker^630^ (AT630), or Nile Red (Dyes); scale bar = 1 µm. (C) Chymotrypsin-like proteasome activity of lysates from octopus was lower than squid or slug tissues (n = 3). (D) Expression of proteasome components from octopus and squid represented as TPM counts for each gene category. Heatmap with tissue-specific expression (brain/arm) of octopus and squid genes part of proteasomal core particle subunit, 26S regulatory subunit, assembly chaperones, and ubiquitin ligases (E); Unfolded protein response (F); and Heat shock proteins (G; n = 3). H**,** The octopus proteasome showed high levels of relaxed selection in contrast to the squid proteasome. Bars show the proportion of proteasomal genes experiencing different kinds of selection (relaxed, neutral, or diversified) relative to a background set of housekeeping genes. Data represented as mean ± s.e.m. *p<0.05, **p<0.01, ***p<0.005 as determined by ANOVA and unpaired Welch’s t-tests.

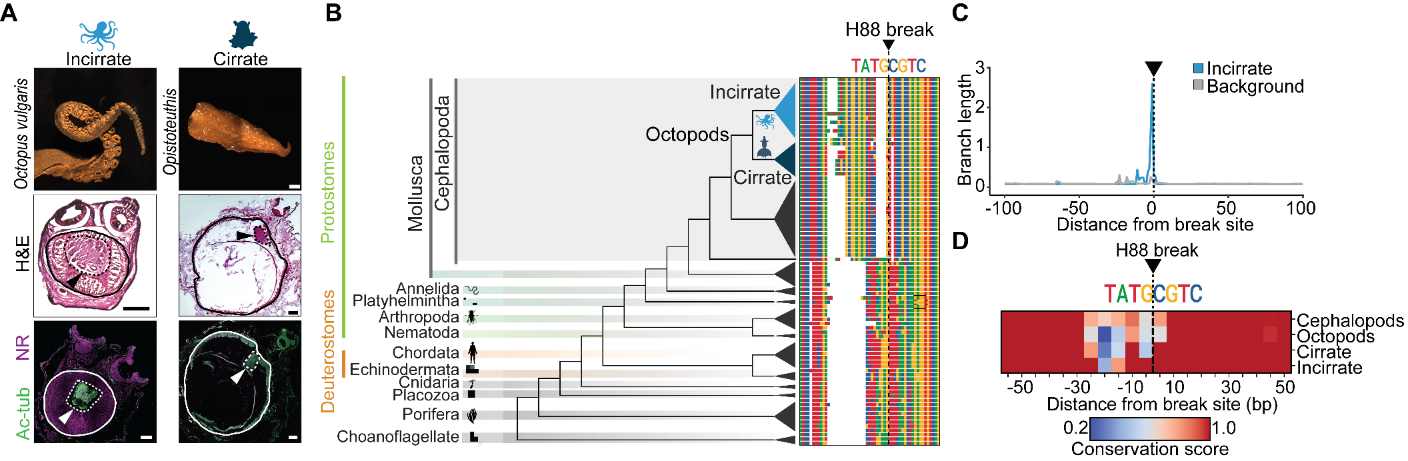

Figure S6: **Lineage-specific evolution of the H88 break site in Incirrate octopuses** (A) Hematoxylin and eosin (H&E) staining and immunohistological labelling using acetylated tubulin antibody of arm cross sections across Incirrate (*Octopus vulgaris*) and Cirrate octopuses (*Opistoteuthis*) reveal an expansion of the nerve cord in Incirrates. Scale bar=500 µm. (B) The octopus H88 break occurs in a site of high conservation across species and is specific to octopuses. Tree shows the evolutionary relationship of species and the aligned consensus rDNA around the break site and is shown without branch lengths for display purposes. (C) Sliding window analysis showing the number of substitutions on the branch leading to Incirrates compared to other cephalopods. (D) Binned conservation scores of rDNA sequences for different cephalopod taxa.

Supplementary Table 1: **Cryo-EM data collection and refinement statistics**

|  | Empty 80S  EMD-48049 | Hybrid 80S  PDB 9EHO  EMD-48061 | 60S (multibody)  EMD-48050 | | 40S (multibody)  EMD-48051 |
| --- | --- | --- | --- | --- | --- |
| **Data collection and processing** | |  |  |  | |
| Magnification | 36,000 | 36,000 | 36,000 | 36,000 | |
| Voltage (kV) | 200 | 200 | 200 | 200 | |
| Electron exposure (e–/Å^2^) | 53 | 53 | 53 | 53 | |
| Defocus range (μm) | -1.5 to -2.5 | -1.5 to -2.5 | -1.5 to -2.5 | -1.5 to -2.5 | |
| Pixel size (Å) | 1.1 | 1.1 | 1.1 | 1.1 | |
| Symmetry imposed | C1 | C1 | C1 | C1 | |
| Initial particle images (no.) | 113,630 | 113,630 | 113,630 | 113,630 | |
| Final particle images (no.) | 30,629 | 15,061 | 69,888 | 69,888 | |
| Map resolution (Å)  FSC threshold | 3.2 0.143 | 3.1 0.143 | 3.1 0.143 | 3.3 0.143 | |
| Map resolution range (Å) | 3.0 to 14.7 | 2.7 to 16.8 | 2.8 to 24.3 | 3.0 to 24.7 | |
| **Refinement** |  |  |  |  | |
| Initial model used (PDB code) |  | 6SGC, ModelAngelo |  |  | |
| Model resolution (Å)  FSC threshold |  | 3.2 0.5 |  |  | |
| Model resolution range (Å) |  | 3.2 to 90.0 |  |  | |
| Map sharpening *B* factor (Å^2^) |  | -51.1 |  |  | |
| Model composition  Non-hydrogen atoms  Protein residues  Nucleotides  Ligands |  | 195,312  10,965  4,969  7 Zn^2+^  205 Mg^2+^  1 SPD |  |  | |
| *B* factors (Å^2^)  Protein  Nucleotide  Ligand |  | 33.83  36.75  16.20 |  |  | |
| R.m.s. deviations  Bond lengths (Å)  Bond angles (°) |  | 0.004  0.774 |  |  | |
| Validation  MolProbity score  Clashscore  Poor rotamers (%) |  | 1.82  8.15  0.01 |  |  | |
| Ramachandran plot  Favored (%)  Allowed (%)  Disallowed (%) |  | 94.50  5.49  0.01 |  |  | |

### Materials and Methods

**Animal sedation and tissue harvesting**

California two-spot octopuses (*O. bimaculoides*) were wild-caught male and female adults (aged 1–2 years, Aquatic Research Consultants) and fed a daily diet of fiddler crabs (*Leptuca pugilator*, Northeast Brine Shrimp, Oak Hill, FL). Adult (aged 6–8 months) hummingbird bobtail squid (*E. berryi*) were laboratory cultured (Marine Biological Laboratory, Woods Hole, MA) and fed daily with grass shrimp (*Palaemonetes pugio*, Aquatic Indicators). All cephalopods were euthanized using established methods with sedation in 15 % ethanol followed by pithing for tissue harvesting. All housed animals were kept under a 12 h–12 h light/dark cycle in filtered natural seawater. Briefly, adult (aged 5–8 months) California sea hares (*A. californica*) were laboratory cultured (National Resource for Aplysia, Miami, FL), anesthetized in 5–10 animal volumes of 1:1 seawater:isotonic MgCl_2_, and dissected upon arrival at Harvard University. Piranhas were euthanized in 100 mg L^-1^ MS-222.

All other animal specimens used for RACE analyses were euthanized, and harvested for tissues upon arrival at Harvard University. Animal protocols were approved by the Harvard University Animal Care and Use Committee (protocol ID 18-05-325 and 18-05-324) with further guidance from the Marine Biological Laboratory Marine Resources Center. Tissues were obtained from the following species and vendors: Caribbean reef octopus (*Octopus briareus*; Pete’s Aquarium; Fishkill, NY); Ruby octopus (*Octopus rubescens*; TONMO); Pyjama squid, Longfin squid, Stumpy cuttlefish, Flamboyant cuttlefish (*Sepioloidea lineolata*, *Doryteuthis pealeii*, *Sepia bandensis*, *Metasepia pfefferi*; Marine Biological Laboratory Woods Hole, MA); Flame scallop, Periwinkle snail (*Ctenoides scaber, Littorina littorea;* Live Aquaria, Rhinelander, WI); Blue mussel, Chiton (*Mytilus edulis, Chaetopleura apiculate;* Gulf of Maine Inc.; Pembroke, ME); Leech, Earthworm (*Hirudo medicinalis, Lumbricus terrestris,* Carolina Biological Supply (Burlington, NC), Red-bellied piranha (*Pygocentrus nattereri,* AquaScapeOnline (Belleville, NJ).

**RNA extraction and gel electrophoresis**

Tissue was flash-frozen in liquid nitrogen immediately after collection and ground into a fine powder using a mortar and pestle. RNA extraction under denaturing conditions was performed by mixing frozen tissue powder with 10× volume TRIzol (Invitrogen) and isolating the RNA aqueous phase according to the manufacturer’s protocol. RNA was further purified using an RNA Clean & Concentrator kit (Zymo). RNA extraction under non-denaturing conditions was performed via phenol-chloroform purification. Tissue powder was resuspended in 3× volume lysis buffer (50 mM Tris-HCl pH 7.5, 10 mM MgCl_2_, 100 mM 2-mercaptoethanol, 0.5 % NP-40 (v/v)), triturated through an 18-g needle, incubated on ice for 10 min, and nuclei were pelleted by spinning at 11,500 × g for 10 min at 4 °C. Total RNA was then purified from lysates by phenol-chloroform extraction and ethanol precipitation and RNA was resuspended in non-denaturing resuspension buffer (10 mM Tris-HCl pH 7.5, 150 mM NaCl, 5 mM MgCl_2_, 1 mM EDTA-NaOH pH 8). For denaturing RNA gel electrophoresis, RNA was mixed with an equal volume of 2× stop solution (95 % deionized formamide, 20 mM EDTA-NaOH pH 8), incubated at 95 °C for 5 min, and analyzed on 1 % (w/v) agarose gels prestained with 2 % (w/v) ethidium bromide. For non-denaturing RNA gel electrophoresis, RNA was mixed with 6× non-denaturing loading dye (40 % w/v sucrose, with bromophenol blue) and analyzed on 1 % (w/v) agarose gels prestained with 2 % (w/v) ethidium bromide.

**RACE-PCR**

For 3′ RACE-PCR analysis, RNAs were polyadenylated by incubating 5 µg of total RNA in a 25 µL reaction containing 10 mM ATP, 40 U mL^-1^ yeast poly-A polymerase (Thermo Fisher), 1× Poly(A) Polymerase Reaction Buffer (Thermo Fisher), and 25 mM MnCl_2_ for 10 min at 37 °C. RNA was isolated using a RNA Clean & Concentrator kit (Zymo) and cDNA was generated using Marathon RT (Addgene #109029) and a 3′-RACE-anchor primer (5′-GACCACGCGTATCGATGTCGACTTTTTTTTTTTTTTT-3′) under standard conditions^6^. PCR was subsequently performed using Q5 polymerase (NEB) and H88-3′-RACE-F (5′- TGAAAAACCGCCACTTCCGTAGTTTC-3′) and 3′ RACE-R (5′- GACCACGCGTATCGATGTCGAC-3′) primers under standard conditions. Products were gel extracted and used as template for a second round of PCR amplification. PCR products were inserted into pMiniT 2.0 by PCR cloning (NEB) and the 3′ end of the rRNA was identified by Sanger sequencing. For 5′ RACE-PCR analysis, 3 μg total RNA was annealed with random hexamers as the reverse transcriptase (RT) primer. RT and template switching was performed using the Template Switching RT Enzyme Mix (NEB) and a template switching primer (5′-GCTAATCATTGCAAGCAGTGGTATCAACGCAGAGTACATrGrGrG-3′) according to manufacturer’s protocol. PCR was subsequently performed using Q5 polymerase and TSO-R (5′-CATTGCAAGCAGTGGTATCAAC-3′) and the H88-5′-RACE-R (5′- GCGAGCGCGCAAACCGTTGC-3′) primers. PCR products were gel extracted, re-amplified, cloned into vectors, and sequenced as described above.

**Ribosome purification**

Ribosomes were isolated from flash-frozen ground tissue powder from *O. bimaculoides* arms*, E. berryi* arms, and *Aplysia* parapodia. Tissue powder was homogenized in lysis buffer (320 mM sucrose, 50 mM Tris-HCl pH 7.5, 100 mM KCl, 10 mM MgCl_2_, 10 mM DTT, 10 U mL^-1^ murine RNase inhibitor (NEB), 0.1 % (v/v) Triton-X 100) and triturated sequentially through a 16-g, 18-g, and 21-g needle. Insoluble material was pelleted by spinning 13,500 × g for 10 min at 4 °C. Lysates were loaded onto a 50 % (w/v) sucrose cushion made in lysis buffer without detergent for 18 h at 4 °C at 30,000 rpm in a Type 70.1 Ti rotor. Pelleted ribosomes were resuspended in storage buffer (6 % (w/v) sucrose, 100 mM KOAc, 5 mM MgCl_2_, 30 mM HEPES-KOH pH 7.5, 2 mM DTT), and further purified by centrifugation through S-400 resin for 2 min at 4 °C at 700 × g^7^.

**Cryo-EM structure determination**

80S ribosomes were isolated for Cryo-EM from octopus arm tissue as described above, but with 10 μM emetine added to the lysis and sucrose cushion buffers. 3 µL of purified octopus monosomes at ~ 140 nM were applied to glow-discharged R2/2 Cu 300 mesh grids (Quantifoil) covered with a 5 nm layer of continuous carbon and vitrified using a Vitrobot Mark IV (Thermo Fisher Scientific) set at 4 °C and 100 % humidity with a wait time of 30 sec, a blot time of 3 sec, and a blot force of +8. Grids were imaged on a Talos Arctica TEM (Thermo Fisher Scientific) operating at 200 kV equipped with a K3 summit direct electron detector (Gatan) in counting mode at a nominal magnification of 36,000× corresponding to a calibrated pixel size of 1.1 Å. Semi-automated data collection was performed using SerialEM v4.0.5. A total exposure time of 4.5 s, corresponding to a total dose of 53 electrons/Å^2^, was fractionated over 50 frames. The defocus range was -1.5 to -2.5 µm.

988 movies were subjected to motion correction and CTF estimation using CTFFIND-4.1^8^ in RELION-3.1^9^, followed by template-free Laplacian-of-Gaussian (LoG)-based automatic particle picking. Following 2D classification, 113,630 particles were subjected to 3D auto-refinement using a 60 Å low-passed initial volume of a rabbit 80S ribosome and 3D classification with 6 classes. 69,888 particles in classes corresponding to 80S ribosomes were subjected to Bayesian polishing and another round of 3D auto-refinement. This dataset was then either directly subjected to multibody refinement of the 60S and 40S ribosomal subunits or to signal subtraction and focused 3D classification without alignments into 6 classes using a mask covering the tRNA binding sites at the ribosomal subunit interface. 15,061 particles corresponding to 80S ribosomes containing hybrid tRNAs and 31,629 particles corresponding to empty 80S ribosomes were separately subjected to a final round of 3D auto-refinement and post-processing in RELION.

The 60S and 40S rabbit ribosomal subunits of PDB 6SGC^10^ were separately fitted as rigid bodies into the maps generated from multibody refinement in ChimeraX v1.3^11^. One round of Phenix v1.20.1 real space refinement^12^ with only rigid body fitting enabled was performed, and the two ribosomal subunits were initially modelled separately in Coot v0.9.6^13^ using the maps generated from multibody refinement. Protein and rRNA sequences were changed to octopus sequences assembled from available annotations and RNA sequencing data (PRJNA658966^14^). Where needed, outputs from ModelAngelo^15,16^ were used to inform rRNA paths. Several rounds of Phenix v1.20.1 real space refine and manual modelling in Coot were performed before the two ribosomal subunits were rigid body fitted into the map of the empty 80S octopus ribosome in ChimeraX v1.3. Cycloheximide was added manually in Coot v.0.9.6 followed by multiple rounds of manual adjustments in Coot and Phenix v.1.20.1 real space refine. The empty 80S octopus ribosome model was then used as a starting model for the hybrid 80S octopus ribosome. The 60S and 40S ribosomal subunits were fitted separately in ChimeraX v1.3, followed by one round of Phenix v.1.20.1 real space refine with only rigid body fitting. tRNAs were added in Coot based on initial models from PDB 6SGC, followed by multiple rounds of manual adjustments in Coot and Phenix 1.20.1 real space refine. Model validations were performed with MolProbity^17^ in Phenix. Structural interpretations and comparisons were done in Coot v0.9.6, PyMOL v2.5.3, and ChimeraX v1.5 or v1.6. Figure panels were made with ChimeraX v1.5 and v1.6. rRNA structural and sequence comparisons with human ribosomes were performed using PDB 6Y57 or 6OLE and NCBI gene entry RNA28SN5.

**DMS probing**

Endogenous ribosomes were purified from human (HEK293T) cells or octopus arm tissue by centrifugation of cell lysate on a sucrose cushion for 20 h at 30,000 rpm using a TLA70.1 rotor. The pelleted ribosomes were resuspended and washed through a high salt cushion (50 % (w/v) sucrose, 50 mM Tris-HCl pH 7.5, 10 mM MgCl_2_, 0.1 % (v/v) Triton-X 100, 500 mM KCl) to remove tRNAs by centrifugation for 20 h at 4 °C at 39,000 rpm in a TLA1001.1 rotor. Purified ribosomes were incubated with vehicle (ethanol) or 0.1% of DMS (Fisher scientific) for 30 min, followed by rRNA extraction using TRIzol (Invitrogen) according to the manufacturer’s protocol. Purified rRNA was reverse transcribed using MarathonRT (in house purified from Addgene #109029), followed by cDNA precipitation and purification. Amplicon sequencing was performed with the following primer pairs tiling H89-H93. Octopus: (5'-AGCTTGACTCTAGTCTGGCACG-3', 5'-ACTGACCCGGTGAGGCG-3'); (5'-GTCAAACGGTAACGCAGGTGT-3', 5'-GCCTCACGATCCTTCTGACCTT-3'); (5'-AATTCACCAAGCGTTGGATTGTTCAC-3', 5'-TGGTGTATGTGCTTGGCTGAGGA-3'); (5'-GCGCCAGAAGCGAGAGC-3', 5'-AGGTTTCTGTCCTCCCTGAGC-3'); (5'-CTGACACCTCCTGCTTAAAACCCA-3', 5'-GCTCACGTTCCCTATTAGTGGGT-3'); (5'-AGATGGTAGCTTCGCCCCATT-3', 5'-TCCCGCGCGCG-3'). Human: (5'-AGCTTGACTCTAGTCTGGCACG-3',5'-ACTGACCCGGTGAGGCG-3'); (5'-GTCAAACGGTAACGCAGGTGT-3', 5'-GCCTCACGATCCTTCTGACCTT-3'); (5'-AATTCACCAAGCGTTGGATTGTTCAC-3', 5'-TGGTGTATGTGCTTGGCTGAGGA-3'); (5'-GCGCCAGAAGCGAGAGC-3', 5'-AGGTTTCTGTCCTCCCTGAGC-3'); (5'-CTGACACCTCCTGCTTAAAACCCA-3', 5'-GCTCACGTTCCCTATTAGTGGGT-3'); (5'-AGATGGTAGCTTCGCCCCATT-3', 5'-TCCCGCGCGCG-3’).

**Ribosome-reconstituted *in vitro* translation**

To generate mutant Renilla luciferase plasmids for translation error experiments, the pFR-CrPV-RLuc plasmid^18^, which contains the cricket paralysis virus (CrPV) IRES followed by Renilla luciferase, was prepared by digestion with PciI and XbaI, followed by two-insert Gibson cloning to introduce the mutated sequence. RNAs for luciferase reporter assays were transcribed using T7 RNA polymerase using a purified PCR product as template^19^.

Endogenous ribosomes were pelleted from rabbit reticulocyte (RRL, Promega) by centrifugation for 4 h at 4 °C at 100,000 rpm in a TLA 100.4 rotor. Ribosome-depleted *in vitro* translation extracts were collected, flash-frozen in single use aliquots, and stored at -80 °C. Before addition to *in vitro* translation extracts, ribosomes were treated with 4 U µL^-1^ micrococcal nuclease (NEB) to digest endogenous ribosome-bound mRNAs. For *in vitro* translation, each reaction contained 55 % (v/v) ribosome-free RRL and 1 μg *in vitro* transcribed mRNA in a final reaction containing 40 nM purified ribosomes, 20 µM amino acids (Promega), and 1.27 U uL^-1^ murine RNase inhibitor (NEB). G418 was added to a final concentration of 70 µg mL^-1^ as indicated. Translation reactions were incubated for 2 h at 30 °C, luciferase assay was performed (GeneCopoeia), and relative luminescence units were measured using a Glomax Multi+ plate reader (Promega). For measuring translation fidelity, *in vitro* translation was performed in paired assays using WT or mutant luciferase mRNA^20^. Point mutations are in the catalytic site of luciferase and therefore amino acid misincorporation is required for luciferase activity. Translation error was quantified by normalizing luciferase activity from *in vitro* translation of mutant to wild-type luciferase mRNA.

**Reconstitution of the H88 break in *E.* coli ribosomes**

Mutant *E. coli* ribosomes were reconstituted using the iSAT system^21^. Briefly, fidelity reporter plasmids for the iSAT system were generated by mutating pk7-Luc^21^ by site-directed mutagenesis to contain a missense mutation at position K529 in firefly luciferase. To introduce the H88 rRNA break into *E. coli* ribosomes, pT7rrn which encodes the *E. coli* rRNA was mutated to add 23S 3′ and 5′ processing stem sequences between 23S rRNA nucleotides 2411 and 2412 by around-the-horn PCR and ligation. 70S ribosomal proteins (TP70) and S150 cell extract were prepared as described previously^22,23^. Each iSAT reaction was performed in a final 10 μL volume and contained 3.3 μL S150 extract, 300 nM TP70 protein, 60 μg mL^-1^ T7 RNA polymerase, 1 μL of 40 % (v/v) PEG-8000 in a final concentration of 8 mM magnesium glutamate, 10 mM ammonium glutamate, 130 mM potassium glutamate, 0.85 mM GTP, 0.85 mM UTP, 0.85 mM CTP, 1.2 mM ATP, 34 μg mL^-1^ folinic acid, 0.171 mg mL^-1^ *E. coli* tRNA, 0.33 mM NAD, 0.27 mM CoA, 4 mM oxalic acid, 1 mM putrescine, 1.5 mM spermidine, 57 mM HEPES, 2 mM amino acids, 37 mM PEP, 4 ng μL^-1^ pk7-Luc reporter wild-type or fidelity mutant plasmid, and 13.3 ng μL^-1^ pT7rrn wild-type or H88 mutant plasmid. iSAT reactions were incubated for 8 h at 37 °C to allow for ribosome assembly and reporter expression. Either 2 μL (for wild-type pk7-Luc) or 10 μL (for fidelity mutant pk7-Luc) of the iSAT reaction were added to 30 μL ONE-Glo Luciferase assay reagent (Promega) on ice, and measured on a plate reader for 20 min at 22 °C. The maximum luminosity of this measurement was used to calculate Luc abundance in each reaction. Subsequently, RNA was isolated from iSAT reactions using a RNeasy RNA extraction kit (Qiagen). For RNA gel electrophoresis, 1 μg RNA was analyzed on a 1 % agarose gel made with 1× TAE buffer, 1 % bleach and 1× SYBRsafe stain.

**A-site tRNA filter binding**

BsT-TyrCUA, a plasmid containing a T7 promoter followed by ß-*stearothermophilus* tyrCUA gene, was used as template for synthesis of the amber suppressor tRNA^24^. The plasmids containing modified anticodon sequences were prepared by around-the-horn PCR and ligation using a shared Tyr-ATH-R primer (5′-AGTCCGCCGCGTTTAGCCACT-3′) and one of the following primers: Tyr-UUA-ATH-F (5′-TTAAATCCGCTCCCTTTGGGTTC-3′), Tyr-UCA-ATH-F (5′-TCAAATCCGCTCCCTTTGGGTTC-3′), Tyr-UGA-ATH-F (5′-TGAAATCCGCTCCCTTTGGGTTC-3′), Tyr-UAA-ATH-F (5′-TAAAATCCGCTCCCTTTGGGTTC-3′). Suppressor tRNA templates for *in vitro* transcription were prepared by digestion of the WT and variant BsT-TyrCUA plasmids with bstNI. T7 *in vitro* transcription of radiolabeled suppressor tRNA was performed as previously described by using final concentrations of 50 μM UTP and 0.4 µCi µL^-1^ [ɑ-^32^P]-UTP (Perkin Elmer) for radiolabeling^25^.

A plasmid containing the *ActB* 5′ UTR followed by AUG-UAG codons was prepared by amplification of the UTR sequence from HEK293T cDNA and Gibson cloning into pcDNA4 Rluc^26^. The template for *in vitro* transcription of start-stop mRNAs (the *ActB* 5′ UTR followed by AUG and stop or near-cognate codons UAG, UGA, or UAA) was prepared by PCR of pcDNA4 *ActB* 5′ UTR Rluc using a shared T7-ActB-F primer (5′-TAATACGACTCACTATAGGG-3′) and one of the following primers to generate different second codons: T7-stop-UAG-R (5′-ACGTGCCTACATGGTGAGCTGGGGGCGGGTGTGG-3′), T7-stop-UAA-R(5′-ACGTGCTTACATGGTGAGCTGGGGGCGGGTGTGG-3′), T7-stop-UGA-R (5′- ACGTGCTCACATGGTGAGCTGGGGGCGGGTGTGG-3′). *Diphtheria toxin (dtA)* mRNA was transcribed by T7 in vitro transcription using a template synthesized by Integrated DNA Technologies. The concentration of *dtA* mRNA used for binding assays was determined by assessing inhibition of protein synthesis and enrichment of radiolabeled A-site tRNA binding to the ribosome. Recombinant tyrosyl-tRNA synthetase (TyrRS) was expressed and purified from *Escherichia coli* as previously described^25^.

To charge the suppressor tRNAs, 50 µM radiolabeled suppressor tRNA, 2.5 µM TyrRS, and 1 mM 4-Benzoyl-phenylalanine (ChemImpex) were incubated in a final reaction containing 50 mM HEPES-KOH pH 7.4, 100 mM KOAc, and 1 mM MgOAc_2_ for 10 min at 30 °C^27^. To prepare the vacuum apparatus for tRNA binding, nitrocellulose membranes (Prometheus) and Hybond N+ membranes (Cytiva) were pre-soaked in water for 15 min, then in cold RB buffer (50 mM HEPES-KOH pH 7.5, 250 mM KCl, 70 mM NH_4_Cl, 10 mM MgCl_2_, 1 mM DTT) for 30 min. Membranes were installed in a Bio-Dot apparatus with the nitrocellulose membrane on top, excess liquid was removed by vacuum, and wells were washed two times with 200 µL cold RB Buffer. Each well was washed again immediately prior to loading sample. For measuring the on-rate of A-site tRNA binding, 15.5 µL reactions containing 10.5 µL of ribosome-depleted RRL, 4 µL of 300 nM MNase-treated ribosomes, and 1 µL of 1 mM amino acids were incubated at 30 °C for 10 min to allow for ribosome run-off of endogenous mRNA fragments. To inhibit eEF2, the reaction was supplemented with 1 µL of 20 ng µL^-1^ *dtA* mRNA for 10 min, then 1 µL of 600 μM NAD+ and 1 µL of 1 μg µL^-1^ start-stop mRNA was added to the reaction and incubated for 10 min to assemble stalled ribosome-mRNA complexes with a stop codon in the unoccupied A-site of ribosomes. To start A-site tRNA binding, 2 µL of the charged suppressor tRNA was added to each reaction. At each indicated timepoint, 2 µL of the reaction was collected and added to pre-cooled tubes containing 48 µL RB Buffer on ice, and immediately filtered by centrifugation through S-400 resin for 1 min at 4 °C at 700 × g^7^ to isolate stalled ribosome-mRNA complexes. Samples were diluted to 200 µL final volume with RB buffer and immediately filtered through a nitrocellulose membrane in a 96-well Bio-Dot vacuum apparatus. Each well was washed three times with 200 µL of RB buffer, the membrane was air dried for 10 min and imaged using a phosphorimager. For near-cognate binding assays, reactions were assembled as described above but in 5 µL final reaction volume, and used for filter binding after 20 min of incubation. To quantify the binding of near-cognate tRNAs, the intensity values from reactions programmed with near-cognate tRNAs were normalized to the reactions set up in parallel with cognate tRNAs for each ribosome.

**Protein aggregate isolation and quantification**

For measuring native protein aggregation, frozen tissue powder was resuspended in 15× volume of fractionation buffer (20 mM Tris-HCl pH 8, 1 mM EDTA-KOH pH 8, 0.33 M sucrose)^28^ and centrifuged for 5 min at 4 °C at 2000 × g. The supernatant was centrifuged for 10 min at 4 °C at 6000 × g, and the resulting supernatant was taken as a total protein sample. 1.5 mg of total protein was centrifuged for 90 min at 4°C at 66,000 rpm in a TLA-110 rotor. The pellet, which contains microsomal membranes and aggregated proteins, was resuspended in 3 mL of fractionation buffer with 0.3 % (v/v) Triton-X 100, incubated for 16 h at 4 °C with continuous shaking, and centrifuged for 60 min at 4°C at 50,000 rpm in a TLA-110 rotor to further fractionate away membranes. The aggregate pellet was resuspended in fractionation buffer with 0.3 % (v/v) Triton-X 100 and centrifuged for 20 min at 4 °C at 16,000 × g. The pellet was resuspended in 150 µL of fractionation buffer with 0.3 % (v/v) Triton-X 100 and resuspended by three rounds of sonication at 25 % power for 10 s using a Q700 sonicator (QSonica). Total and aggregate protein was separated by SDS-PAGE, stained with Coomassie Blue, and gel lane intensity was calculated using ImageLab.

To generate HA-tagged Firefly luciferase plasmids for aggregation assays, pcDNA4 *PSMB6* 5' UTR FLuc^29^ was used to generate pcDNA4 CrPV IRES HA-Fluc by Gibson cloning. RNAs containing inosine were transcribed as above but with 2.9 mM ATP and 0.1 mM ITP. To measure Firefly luciferase aggregation, *in vitro* translation reactions were assembled as described above but using HA-tagged Firefly luciferase mRNA and incubated for 3 h at 30 °C, a timepoint chosen to obtain equivalent total levels of luciferase protein synthesized by all ribosomes. The reaction was fractionated into soluble and aggregate fractions by centrifugation for 40 min at 4 °C at 91,000 rpm in a TLA 100.4 rotor. Pelleted aggregated protein was resuspended in 10 µL of 8 M urea and incubated at 23 °C for 1 h with mixing by pipetting. Soluble and aggregate protein was analyzed by immunoblot with an anti-firefly luciferase antibody (Proteintech, 27986-AP-1). Gel band intensity was calculated using ImageLab (BioRad).

**Proteasome activity assays**

Frozen tissue powder was resuspended in 3× volume ice cold PBS-E (PBS with 5mM EDTA) and homogenized by sonication at 25 % power for 6 sec. Lysate was centrifuged for 5 min at 4 °C at 13,000 × g and the supernatant diluted to a final concentration of 1.3 mg mL^-1^ protein concentration in cold PBS-E. 50 µL of Proteasome-Glo reagent (Promega) was added, samples were incubated for 1 h at 23 °C, and proteasome activity was measured as luminescent signal intensity in a 96-well plate reader (Promega).

**RNA sequencing and bioinformatics**

Tissue samples for RNA-seq were collected from animals in RNAlater and flash frozen liquid nitrogen. RNA extraction, library prep and sequencing using HiSeq Rapid Run were performed by Azenta Life Sciences. Reads were quality filtered and trimmed using Trim Galore. Reference transcriptomes were de novo assembled using Trinity^30^ and open reading frames were determined using Transdecoder. Reads were pseudo-aligned and transcript abundance was estimated using Kallisto using the de novo transcriptome assembly as the reference^31^ . Annotation was performed using DIAMOND^32^. Sequencing data generated from this study are available at the NCBI Sequence Read Archive under BioProject PRJNA1199547 and GEO Accession number GSE284564.

**rDNA phylogenomics**

To understand how the break site evolved in cephalopods, a blast search was performed against the non-redundant nucleotide database using a query sequence that was 50 bases up and downstream of the break site from *Octopus bimaculoides*. The full sequences from these queries from Genbank were downloaded using the read.genbank function in R. These sequences were aligned with MAFFT v7.490^33^ using the L-INS-I algorithm. A constrained and dated phylogenetic tree was built using IQtree. To constrain the topology of the tree based on prior studies, the following constraint tree was used: “(((((Eledone cirrhosa,Octopus sinensis),Grimpoteuthis sp.),Vampyroteuthis infernalis),(Euprymna morsei,Sepietta oweniana)),Nautilus pompilius);”. To date the tree, dates were included from a recent phylogenomic study on mollusks^34^. The best fitting model of sequence evolution was found with ModelFinder^35^ implemented in IQtree, using 1000 ultrafast bootstraps to get support measures for nodes^36^.To understand whether the break site was a lineage specific innovation in Incirrates, a sliding window analysis using 10 nucleotide windows was performed that calculated the branch length of the branch leading to Incirrate octopuses along with the average branch length across the entire cephalopod tree.

To understand conservation in the ribosome outside of cephalopods, we collected ribosomal RNA sequences across a diverse range of animals with a focus on high-quality long-read genomes but including a few short-read genomes of phylogenetically important organisms. Ribosomal RNA sequences were found in genomes by using blastn^37^ with an e-value cut-off of 1×10^-5^ using the *Octopus bimaculoides* genomic rDNA around the break site as a query. Sequences shorter than 300 nucleotides were removed and surviving sequences were clustered using cd-hit^38^ (clustering threshold of 1). Since rDNA exists as tandem repeats, multiple rDNA sequences were retrieved for each genome. To deal with these repeats, sequences were first aligned using MAFFT v7.490^33^ using the L-INS-I algorithm, then poorly aligning sequences were removed with trimAl^39^. Consensus rDNA sequences were generated for each species using DECIPHER and used for downstream analysis. These sequences were displayed with the cephalopod tree produced above along with branches representing major animal lineages.

**Inference of relaxed or intensified selection**

To understand whether the H88 break had downstream effects on the evolution of other components of the protein synthesis and degradation pathway, rates of relaxed or intensified selection were compared. To identify relevant genes, human proteins that were annotated with eggnogmapper^40^ were used. Homologs across molluscan genomes were found by identifying the reciprocal best blast hit of the human proteins. Blast hits were required to survive an e-value cut-off of 10^-10^. Proteins were aligned using MAFFT (L-INS-I) and protein alignments were used to generate codon alignments with PAL2NAL^41^. Relaxed selection in octopus and squid lineages was estimated with RELAX^42^ on both untrimmed and trimmed codon alignments (trimmed with the -gappyout flag in TrimAl). Based on the results from RELAX, octopus and squid proteins were categorized as undergoing relaxed, intensified, or no change in selection relative to background. To test for increases in relaxed selection relative to the background gene set, a binary variable was generated with two states: not relaxed (genes that were found to be experiencing no change or intensified selection) and relaxed. Increases in relaxed selection were then estimated using a logistic regression model, and increased relaxed selection was tested on both untrimmed and trimmed codon alignments.

**Immunohistochemistry**

Cephalopod arms which were either freshly dissected from wild caught animals or obtained from museum samples stored in 70 % ethanol were fixed in PFA buffer (PBS with 4 % (v/v) paraformaldehyde) at 4 °C for 16–24 h with rocking. Following fixation, tissues were washed with PBS and stored in 50 % (w/v) sucrose at 4 °C for 48–72 h, embedded in O.C.T (Tissue-Tek, Sakura Finetech) and frozen at -80 °C for at least 24 h. Embedded tissues were sectioned using a cryostat (Leica CM3050S) at 20 μm sections. Sections were dried at room temperature for 30–60 min and stored at -20 °C overnight. After serial washes in PBST buffer (PBS with 0.3 % Triton X-100), samples were blocked for 1 hr in blocking buffer (PBST buffer with 5 % (v/v) normal goat serum, 0.2 % (w/v) bovine serum albumin) and incubated with 50 µL primary antibody solution overnight. For neuronal staining, primary antibody was a mouse monoclonal anti-acetylated tubulin (Sigma, T6793), followed by staining with a goat anti-mouse AlexaFluor^555^ secondary antibody (Invitrogen, A-21422) for 2–4 h at room temperature in blocking buffer. Samples were washed 3–5 times in PBST buffer, mounted in Vectashield with DAPI (Vector Laboratories, H-1200), and imaged with a Zeiss LSM900 confocal. For general tissue histology, sections were stained with hematoxylin and eosin dye (Abcam, ab245880) as per manufacturer’s protocol. Protein aggregates in tissue sections were labelled using Proteostat Aggresome detection kit (Enzo Life Sciences, ENZ-51035), Nile Red staining kit (Abcam, ab228553), or Amytracker630 (Ebba Biotech) as per kit instructions.

***Ex vivo* tissue culture**

Octopus and squid arms were sedated in 15 % ethanol and euthanized as per IACUC protocols. Arms were dissected in sterile artificial sea water (430 mM NaCl, 10 mM KCl, 10 mM CaCl_2_, 50 mM MgCl_2_, 10 mM HEPES, 10 mM glucose) and maintained *ex vivo* for 24 hours in a modified Leibovitz’s L15 medium (ThermoFisher) adjusted to 430 mM NaCl, 10 mM KCl, 10 mM CaCl_2_, 50 mM MgCl_2_, 10 mM HEPES, 10 mM glucose, 4 % FBS, 1 % penicillin/streptomycin, at pH 7.8 and 980 mOSM, at 18 °C. Pharmacological treatments of cultured arms were carried out with 2 mg mL^-1^ G418 (Fisher), 10 µg mL^-1^ S-MG132 (Cayman Chemical Company) or DMSO (vehicle control).

**rRNA break analysis RT-PCR**

*Grimpoteuthis* tissue fragments were collected in 95 % ethanol and stored in 50 % ethanol until it was flash frozen in liquid nitrogen and ground into a fine powder using a mortar and pestle. RNA extraction was performed by mixing frozen tissue powder with 10× volume TRIzol (Invitrogen) and isolating the RNA aqueous phase according to the manufacturer’s protocol. RNA was further purified using an RNA Clean & Concentrator kit (Zymo), treated with DNAse I (Thermo Fisher) and converted into cDNA using a High-Capacity cDNA Reverse Transcription Kit (Thermo Fisher). PCR was performed using Q5 polymerase on the cDNA using primers spanning (5′-AGCTCGCTCGATCTCGATTT-3′, 5′-CCACAAGCCAGTTATCCCTGT-3′) and upstream (5′-AGCAAAAGGGCAAAAGCTCG-3′, 5′-TGAAAGGATCGATGGGCCAC-3′) to the putative break site location. As a control, RNA isolated from *Octopus bimaculoides* tissue was subjected to the same protocol and the cDNA was subjected to PCR using primers spanning (5′-AAGGACAAAGCTCGCTCGAT-3′, 5′- CAAACGCTTAGCCGTCACAA-3′) and upstream (5′-AGCTCGCTCGATCACGATTT-3′, 5′- TTCGGAAAGGATCGATGGGC-3′) to the break site.

**Octopus depth analysis**

To compare where Incirrate and Cirrate octopuses occur in the ocean, research-grade species distribution data for all cephalopods were downloaded from GBIF^43^. These data were categorized as Cirrate or Incirrate based on their Families.

**Data analysis**

All significance tests were justified considering the experimental design and tested for normal distribution and variance, using standard tests for normality. Sample sizes were chosen based on the number of independent experiments required for statistical significance and technical feasibility. All graphs and statistical analyses were performed in GraphPad Prism (GraphPad Software).
